# Energetics of stalk intermediates elucidated from hydrostatic pressure effects on membrane fusion

**DOI:** 10.64898/2026.07.29.741230

**Authors:** Daniel Milshteyn, Jacob R. Winnikoff, Joel E. Morgan, Gonen Golani, Itay Budin

**Affiliations:** Department of Biochemistry & Molecular Biophysics, University of California San Diego, La Jolla, CA, USA; Department of Organismal and Evolutionary Biology, Harvard University, Cambridge, MA, USA; Center for Biotechnology and Interdisciplinary Studies, Rensselaer Polytechnic Institute, Troy, NY, USA; Department of Physics, University of Haifa, 3498838, Haifa, Israel

## Abstract

Membrane fusion is an essential process in cells that requires a balance of lipid composition to establish biophysical properties conducive to topology changes. During fusion, lipids of opposing membranes must invert and overcome energy barriers associated with forming highly curved stalk and pore intermediates. While theoretical work has modelled the effect of lipid intrinsic curvature on stalk formation, quantifying the relationship experimentally has proven to be a challenge due to the inability to vary lipid curvature without concomitantly changing other properties that affect fusion. Here we address this hurdle by using hydrostatic pressure to modulate lipid intrinsic curvature independently of chemical composition. Using high-pressure stopped-flow fluorimetry, we measured rates of calcium-mediated lipid mixing between populations of vesicles, a process that is strongly inhibited by pressure. We correlated mean lipid intrinsic curvature across pressure with lipid mixing rates by incorporating complementary small-angle x-ray scattering measurements for each individual lipid component. This analysis showed that lipid mixing rates, a proxy for hemifusion, across compositional and pressure regimes are determined by changes in lipid spontaneous curvature. Consistent with previous theoretical models, we find a linear relation between lipid intrinsic curvature and the hemifusion stalk formation energy, offering direct experimental support for the stalk hypothesis.

**Significance statement:** Membrane fusion proceeds through a hemifusion stalk intermediate whose formation energy depends on lipid intrinsic curvature, a central prediction of the stalk hypothesis that has lacked direct experimental support. Previous tests relied on changes in lipid composition that affect multiple membrane properties, confounding the contribution of curvature alone. Here we use hydrostatic pressure to tune lipid curvature independently of chemical composition and calibrate its effects with high-pressure SAXS. Hemifusion rates across three lipid compositions and four pressures collapse into a single exponential dependence on mean spontaneous curvature, yielding a linear relation between curvature and energy consistent with continuum elastic theory. This work quantifies how lipid composition tunes fusion kinetics, suggesting that small changes in lipid curvature may strongly affect fusogenicity.

## Introduction

Membrane fusion is a universal biological process that underlies intracellular trafficking, neurotransmitter release, and viral infection (1–3). Both computational modeling and experimental evidence support a common pathway for all fusion events, the stalk hypothesis, with universal intermediate states that involve initial lipid mixing in the formation of a hemifusion stalk (Fig. 1A) and subsequent content mixing through pore formation (1, 4, 5). The main energy barriers in this pathway are dehydration of the opposing membranes and the lipid rearrangements that drive the transition from a lamellar to a nonlamellar topology during stalk formation, followed by the barrier to fusion pore opening as membrane stress builds within the hemifusion diaphragm. While protein machinery such as soluble N-ethylmaleimide-sensitive factor attachment protein receptors (SNAREs), and viral fusion proteins supply the energy and specificity for fusion *in vivo*, the lipid bilayer itself is the substrate that must be deformed, and the energetic cost of that deformation sets the fundamental barriers that these proteins must overcome.

In both stalk and pore formation, lipids transition between highly curved membrane intermediates via transient nonlamellar topologies. The stalk resembles the inverted hexagonal (H_II_) or inverted bicontinuous cubic phase (Q_II_) phase, while the pore rim is positively curved. The formation of the stalk is favored by nonbilayer lipids with an inverted-cone geometric shape, or negative intrinsic curvature (c_0_ < 0), which have a smaller cross-section at the headgroup than in the hydrocarbon region. Lipids with high negative curvature, such as phosphatidylethanolamine (PE), cardiolipin (CL), cholesterol, and diacylglycerol (DAG), have been shown to promote stalk formation and increase lipid mixing rates, while lipids with positive intrinsic curvature (c_0_ > 0), such as lysophosphatidylcholine (LPC), inhibit stalk formation but promote pore formation (1–3, 5). Negatively curved lipids better accommodate the extreme negative splay required at the stalk, reducing the energy cost of forming this intermediate and accelerating hemifusion. Quantitatively, the tilt-splay elastic model describes the energy required to form a hemifusion stalk depending on how much the lipid monolayer must bend away from its preferred curvature (6, 7). Across a biologically relevant curvature range, the relationship between stalk energy and c_0_ is predicted to be linear (7, 8). When combined with Kramers’ rate theory (9), which relates energy barriers to reaction rates, this dependence predicts an exponential dependence of the hemifusion rate with c_0_. Theoretical modeling predicts this relationship (7, 8, 10, 11) but more comprehensive experimental evidence is needed to better quantify and test it.

Experimental approaches have classically used varying lipid compositions to indirectly study the roles of membrane properties in fusion (12–14). However, substitution of lipid species often affects more than one physical parameter, complicating attempts to distinguish energetic contributions of independent mechanical forces in each intermediate state. For example, negatively curved cholesterol increases bending rigidity, while adding positively curved LPC softens it (15–17). Because of this entanglement, estimations from composition-based experiments may not reflect the true contribution of intrinsic curvature to the hemifusion energy barrier. In addition to lipid composition, physical parameters such as temperature (18), osmotic pressure (19), and hydrostatic pressure (20) can be used to manipulate membrane properties. Hydrostatic pressure drives lipids into lower-volume conformations and has a particularly large effect on membrane curvature because phospholipid compressibility is anisotropic (21). The acyl chain volume is more pressure sensitive than the headgroup region, so pressure preferentially compresses the hydrocarbon tails and shifts c_0_ toward less negative values without changing the chemical identity of the lipid. This effect is reflected in the pressure sensitivity of curvature-mediated phase transitions. The dT/dP of the L_α_ to H_II_ transition is ∼0.05 K/bar, whereas the dT/dP for the chain-ordering L_α_ to L_β_ transition is roughly 3-fold smaller (22–24). Therefore, pressure preferentially alters lipid intrinsic curvature c_0_ while exerting comparatively modest effects on membrane fluidity and rigidity.

Here we use a FRET-based lipid mixing assay under a range of hydrostatic pressure conditions to experimentally test the relationship between lipid intrinsic curvature and hemifusion rate. We combine this approach with variations in the PE/PC ratio of vesicle populations and estimates of mean intrinsic curvature using pressure-dependent measurements of individual lipid c_0_ values obtained by high-pressure small-angle X-ray scattering (HPSAXS). Together, we find that hemifusion stalk formation energy is linearly dependent on the mean intrinsic curvature across lipid components, supporting predictions arising from the stalk hypothesis. In addition, simulations of stalk structures across a range of parameters show that the experimental results are consistent with theoretical calculations when relatively soft membrane bending moduli are assumed.

## Methods

### Lipid mixing fusion assay

Membrane fusion kinetics were assessed using a FRET-based lipid mixing assay, in which fusion is reported by fluorescence dequenching upon dilution of a self-quenching donor-acceptor pair. Chloroform stocks of POPC, POPS, and DOPE (Avanti Research) were combined at the desired molar ratios, and the solvent was removed by drying under nitrogen for 10 minutes followed by high vacuum for at least 1 hour. Resulting lipid films were hydrated with Buffer A (100 mM NaCl, 5 mM Na^+^ HEPES, 0.1 mM EDTA, pH 7.4) to obtain 400 µM lipid (2X) and tumbled for 1 h. Unilamellar vesicles of approximately 100 nm diameter were produced by four freeze-thaw cycles followed by extrusion through a 100 nm polycarbonate membrane 21 times. One population of fluorescence-quenched liposomes, containing lissamine rhodamine B labelled DOPE (Rh-DOPE, 2 mol%) and N-(7-Nitrobenz-2-Oxa-1,3-Diazol-4-yl labelled DOPE (NBD-DOPE, 2 mol%) with DOPE:POPC:POPS (32:32:32 mol%), DOPE:POPC:POPS (42.6:21.4:32 mol%), or DOPE:POPC:POPS (21.4:42.6:32 mol%), were mixed with a population of liposomes lacking fluorophores with DOPE:POPC:POPS (33.3:33.3:33.3 mol%), DOPE:POPC:POPS (44.6:23.4:32 mol%), or DOPE:POPC:POPS (23.4:44.6:32 mol%).

The emission of NBD-PE dequenching was measured following the addition of 20 mM CaCl_2_ (2X) in Buffer A, for a final reaction concentration of 200 µM lipid and 10 mM CaCl_2_, using a High-Pressure Stopped Flow System (TgK Scientific) equipped with a Hi-Tech UV–vis spectrophotometer at the Rensselaer CBIS Analytical Biochemistry Core Facility. NBD was excited at 460 nm and its emission monitored through a FITC bandpass filter (513-556 nm), and temperature was maintained at 37 °C using an integral thermostated jacket. Individual runs were carried out with hydrostatic pressures set from 1-750 bar, fluorescence intensity was recorded for 5 min to allow reactions to reach equilibrium. Fluorescence of a fully dequenched control, with 1 mol% NBD-DOPE and Rh-DOPE in DOPE:POPC:POPS (32.65: 32.65: 32.65), DOPE:POPC:POPS (43.6: 22.4: 32), or DOPE:POPC:POPS (22.4: 43.6: 32) was recorded to permit normalization and determination of total % fused vesicles.

### SAXS sample preparation

Lipid stocks of di-oleoyl phosphatidylethanolamine (DOPE), 1-palmitoyl-2-oleoyl phosphatidylcholine (POPC), or 1-palmitoyl-2-oleoyl phosphatidylserine (POPS) (Avanti Research) in chloroform were used to prepare lipid mixtures with a total mass of 5 mg lipid. The solvent was dried under a stream of nitrogen for 5 minutes and high vacuum for 1 hour. Lipids were resuspended in 100 µL cyclohexane and 50 µL aliquots were dispensed into 1.5 mm inner diameter capillary melting point tubes (Kimble Chase 34505-99), flash-frozen at -80 °C, and lyophilized for 1 hour to yield a lyophilized powder. Rehydration was carried out by adding 10 µL of vacuum-degassed MilliQ water, mixing with a 20 µL Wiretrol II microdispenser (Drummond Scientific), followed by five freeze-thaw cycles using an ethanol/dry ice bath. Between cycles, the suspension was homogenized by repeated pipetting with a 20 µL Wiretrol II microdispenser within the capillary tube. The resulting lipid suspension (50% w/v) was consolidated to the tube bottom by centrifugation at 500 RCF for 5 minutes at 4 °C before being scored and cut to fit a sample holder. An additional 5 µL of degassed water was added on top of the suspension to ensure complete hydration of the sample. The capillary was then sealed with a plug of high vacuum grease (Dow Corning) introduced through a 22G × 100 mm needle (Air-Tite), which also served as a pressure-transmitting cap. The final lipid concentration was 250 mg/mL.

### SAXS data acquisition and analysis

Collection of HPSAXS data was performed at beamline 7A in the Cornell High Energy Synchrotron Source (CHESS) using a custom-built temperature-controlled pressure cell (25). Data was acquired using a photon energy of 14 keV, 1s exposures using a flux of 1.8-3.6x10^11^ photons/s, a spot size of 200 x 250 µm (W x H), with a collection range of 0.05 nm^-1^ ≤ q ≤ 7 nm^-1^ using an EIGER 4M detector (Dectris) within the beamline vacuum. After testing each sample for radiation sensitivity with five initial exposures, pressure sweeps were performed from lowest to highest and back with triplicate exposures for each pressure condition and 60s of equilibration per 100 bar of pressure change. No background subtraction was used following assessment of low background of a glass capillary with degassed water. SAXS images were azimuthally integrated using BIOXTAS RAW software (26).

### Measurement of phospholipid monolayer curvature

Hosted mixtures at molar ratios of 10 mol%:90 mol% or 20 mol%:80 mol% guest:host with 12% w/w 9(Z)-tricosene for HPSAXS were prepared using DOPE as the host lipid, as described above. Neutral-plane c_0_ was estimated by using a constant distance correction of 0.394 nm, derived from global fitting of the DOPE H_II_ scattering profile (27). Total c_0_ was calculated from peak spacing as previously described (28), and the guest lipid c_0_ was extracted from a weighted average of the total curvature of the hosted system.

### Hemifusion stalk calculations

The elastic energy of the hemifusion stalk was calculated using the continuum elastic theory of lipid monolayers as described previously (8, 29, 30). In this framework, membrane deformations are described in terms of lipid splay, saddle splay, and tilt degrees of freedom, and the equilibrium stalk structure is obtained by numerical minimization of the total membrane elastic energy. Full details of the theoretical formalism, elastic energy functional, numerical implementation, and minimization procedures are provided in these previous works. Briefly, the membrane is modeled as two coupled lipid monolayers connected by a membrane mid-plane. Each monolayer is characterized by a local lipid director field and elastic contributions associated with splay, saddle-splay, and lipid tilt deformations. The elastic energy is evaluated over the monolayer surfaces and minimized numerically with respect to both membrane shape and lipid director configuration.

For the calculations presented here, the fusion site was assumed to be axially symmetric and initially flat, and the two membranes were taken to be compositionally and mechanically symmetric. The stalk configuration corresponds to the merger of the two proximal monolayers while the distal monolayers remain separated. Similar to previous continuum descriptions of membrane fusion stalks, the lipids near the stalk undergo strong tilt and splay deformations in order to avoid hydrophobic void formation at the fusion center. In contrast to previous calculations, where the angle between the membrane mid-planes at the stalk center was fixed to 90°, here this angle was allowed to vary freely during the minimization procedure. This additional degree of freedom reduced the magnitude of lipid splay in the stalk region and resulted in lower stalk elastic energies compared to earlier calculations with a fixed-angle constraint. The equilibrium stalk structure and corresponding stalk formation energy were obtained by minimizing the total elastic energy relative to the undeformed membrane state using an open-source code available at https://github.com/GonenGolani/Fusion_Solver.

## Results

### Hydrostatic pressure inhibits hemifusion

To characterize the effects of hydrostatic pressure on fusion, we employed a model fusion reaction mediated by the interactions between Ca^2+^ and phosphatidylserine (PS) lipids on opposing membranes (31). Ca^2+^ and PS form a 1:2 complex (32), which has been proposed to specifically mediate membrane fusion reactions by bringing together opposing lamellae. While simplified compared to those employing protein-mediated fusogens, the assay relies solely on lipid components and thus prevents complications from any potential pressure effects on protein conformations and dynamics. It also is compatible with the large sample volumes required for high-pressure stopped flow fluorometry.

We measured Ca^2+^-mediated lipid mixing of large unilamellar vesicles (LUVs) composed of equimolar DOPE:POPC:POPS (33:33:33). We read out lipid mixing using a FRET-based assay with a high-pressure stopped flow system (Fig. 1B), in which two solutions are pressurized and mixed isobarically. In our assays, vesicles labeled with 2 mol% NBD-PE and 2 mol% Rh-PE were rapidly mixed with a solution of unlabeled vesicles and 10 mM CaCl_2_ (final concentration) at 37°C. The increase in NBD fluorescence upon FRET dequenching reported on lipid mixing, or hemifusion, between the two vesicle populations. Fluorescence intensities were normalized to a fully dequenched control to determine the total percentage of fused vesicles. At ambient pressure (1 bar), mixing of equimolar vesicles led to rapid lipid mixing, reaching approximately 20% lipid mixing within 75 s (Fig. 1C).

Increasing hydrostatic pressure progressively reduced both the rate and extent of lipid mixing. At 250 bar, the same vesicles reached roughly 15% lipid mixing, while at 750 bar the signal plateaued below 5%. Rate constants extracted from first-order exponential fits decreased from 0.186 ± 0.009 s^−1^ at 1 bar to 0.047 ± 0.001 s^−1^ at 500 bar (Fig. 1D). On a semilogarithmic scale, ln(k) decreased linearly with pressure, consistent with an Arrhenius-like process. Thus, increasing pressures dramatically reduce rates of calcium-mediated fusion.

**Figure 1:**
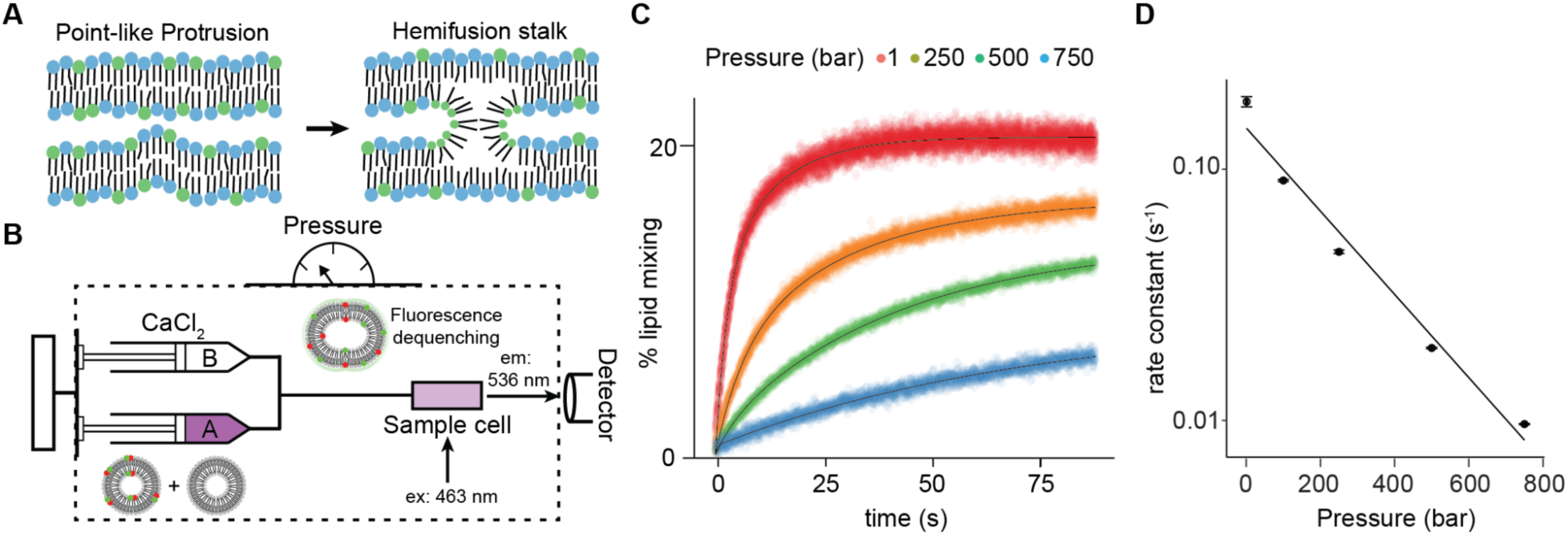
Hydrostatic pressure inhibits lipid mixing during calcium-mediated vesicle fusion. **A**) Schematic of the classical hemifusion pathway. Two bilayers in apposition start at a point-like protrusion evolves into a hemifusion stalk which can be initiated by fusion proteins, the presence of calcium, or osmotic pressure. Stalk formation is supported by the presence of negative curvature lipids, labelled with green headgroups while bilayer-supporting lipids are labelled blue. The hemifusion stalk is a membrane topology characterized by high negative curvature which resembles nonbilayer HII mesophases. **B**) Schematic of high-pressure stopped-flow fluorometry to monitor hemifusion rates at varying pressures. FRET dequenching occurs when vesicles, labelled with 2 mol% NBD-PE and Rh-PE, fuse with unlabeled ones upon the addition of 10 mM calcium. **C**) Lipid mixing assay comparing the hemifusion of liposomes with DOPE:POPC:POPS (33:33:33) in the presence of 10 mM CaCl2 at 37 °C under pressures of 1-750 bar pressure. Averaged intensities are normalized to values for mixed liposomes. Error bars indicate SEM from n=3 replicate runs. **D**) Rate constants from first order exponential fits for lipid mixing of liposomes with DOPE:POPC:POPS (33:33:33 mol%) under pressures of 1-750 bar pressure. Rates decrease with increases in pressure in an exponential relationship. Error bars indicate SEM from n=3 replicate runs.

### Nonbilayer lipid content buffers against pressure inhibition of hemifusion

We hypothesized that the strong inhibition of hemifusion by pressure could result from its effect on lipid intrinsic curvature (c_0_). To examine how lipid composition modulates the pressure sensitivity of fusion, we measured lipid mixing kinetics at two additional DOPE:POPC:POPS ratios — 44:23:32 (high PE) and 23:44:32 (low PE), alongside the equimolar composition, across pressures from 1 to 1000 bar (Fig. 2A). Because DOPE has a strongly negative c_0_ while POPC is approximately cylindrical, increasing the PE:PC ratio makes the mean curvature more negative, which is expected to lower the hemifusion barrier. Consistent with this hypothesis, higher PE content increased fusion rates and capacity across all pressures.

At ambient pressure, the high PE vesicles showed fast lipid mixing approaching 60% hemifusion, while the low PE vesicles reached a plateau at roughly 20%. At elevated pressure, the high PE composition maintained efficient lipid mixing even at 1000 bar, whereas the low PE composition showed near-complete inhibition of fusion by 750 bar. The equimolar composition showed intermediate behavior. Rate constants from first-order exponential fits (Fig. 2B) spanned nearly two orders of magnitude across compositions and pressures, from 0.72 s^−1^ (high PE, 1 bar) to 0.006 s^−1^ (low PE, 750 bar). These results indicate that PE content effectively buffers the membrane against pressure-induced inhibition. On a semilogarithmic scale, ln(k) decreased linearly with pressure at each composition, with similar slopes across the three PE:PC ratios, suggesting that the effect of pressure on the hemifusion barrier is comparable regardless of the membrane lipid composition.

**Figure 2:**
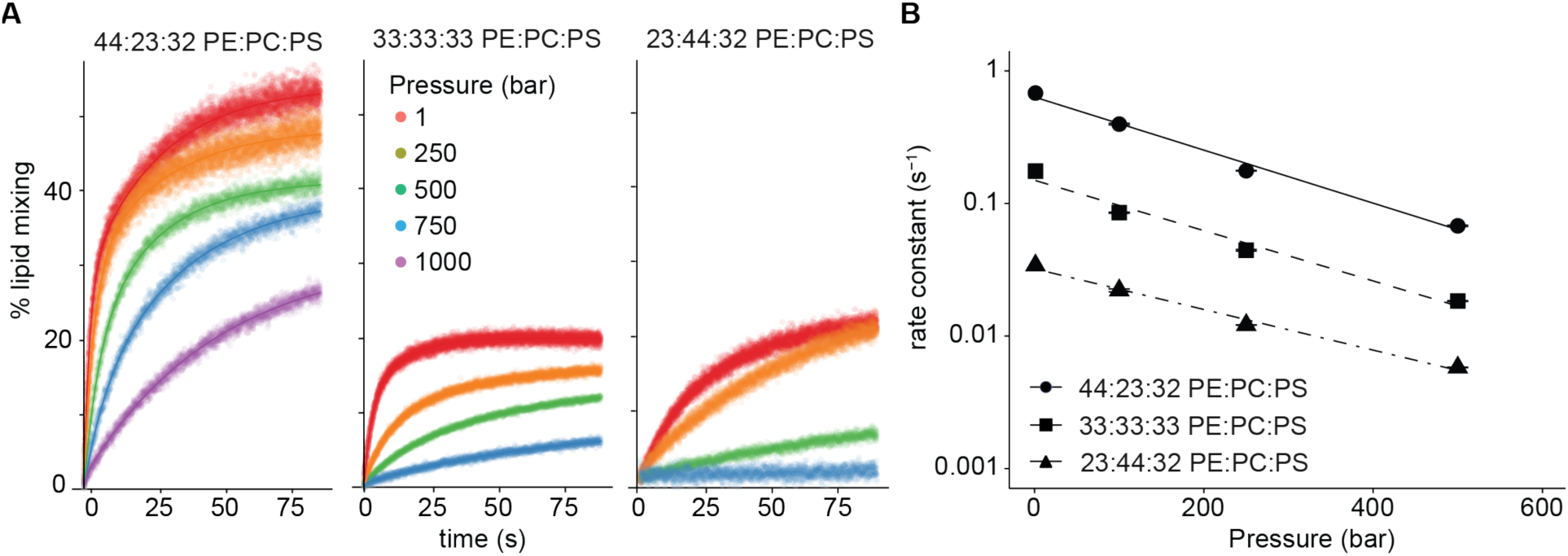
Increased PE content buffers against high-pressure inhibition of hemifusion. **A**) Lipid mixing assay of liposomes with different DOPE:POPC:POPS ratios at 1-1000 bar pressure and 37 °C. High PE (44:23:32 mol% DOPE:POPC:POPS) results in increased lipid mixing rates and capacity, while low PE (23:44:32 mol% DOPE:POPC:POPS) results in inhibited lipid mixing failing by 750 bar pressure. Equimolar (33:23:33 mol% DOPE:POPC:POPS) mixtures showed intermediate lipid mixing to the high and low PE compositions. Averaged intensities are normalized to values for mixed liposomes. Error bars indicate SEM from n=3 replicate runs. **B**) Rate constants from first order exponential fits for lipid mixing of liposomes with 44:23:32 mol%, 33:33:33 mol%, or 23:44:32 mol% DOPE:POPC:POPS under pressures of 1-500 bar pressure. Error bars indicate SEM from n=3 replicate runs.

### Hydrostatic pressure reduces negative intrinsic curvature of phospholipids

To quantitatively assess the role of c_0_ in measured hemifusion rates, we performed HPSAXS on fully hydrated, tricosene-saturated lipid dispersions of DOPE, POPC, and POPS (the latter two hosted at 20 mol% in 80 mol% DOPE) at 35 °C across pressures from 1 to 800 bar (Fig. 3A). In each case, the lipids adopted the H_II_ phase, and the spontaneous curvature was estimated from the first-order Bragg peak position. Addition of POPC or POPS to the DOPE host system shifted the first-order peak to lower q values (Fig. 3B), reflecting their less negative curvature contributions that increase the H_II_ lattice spacing. With increasing pressure, the first-order peak of all samples shifted progressively to lower q, indicating an increase in H_II_ cell size and thus a decrease in negative curvature (Fig. 3C).

Quantification of c_0_ across the pressure range revealed that all three lipid species became less negatively curved with increasing pressure (Fig. 2D). DOPE maintained strongly negative curvature throughout, decreasing from −0.385 nm^−1^ at 1 bar to −0.366 nm^−1^ at 800 bar. POPC shifted from −0.075 nm^−1^ toward near-zero values with a cylindrical geometry. POPS exhibited a positive c_0_ that increased further under pressure, moving from 0.043 nm^−1^ at 1 bar to 0.079 nm^−1^ at 800 bar. All three lipids showed linear pressure dependencies with similar slopes (dc_0_/dP).

**Figure 3:**
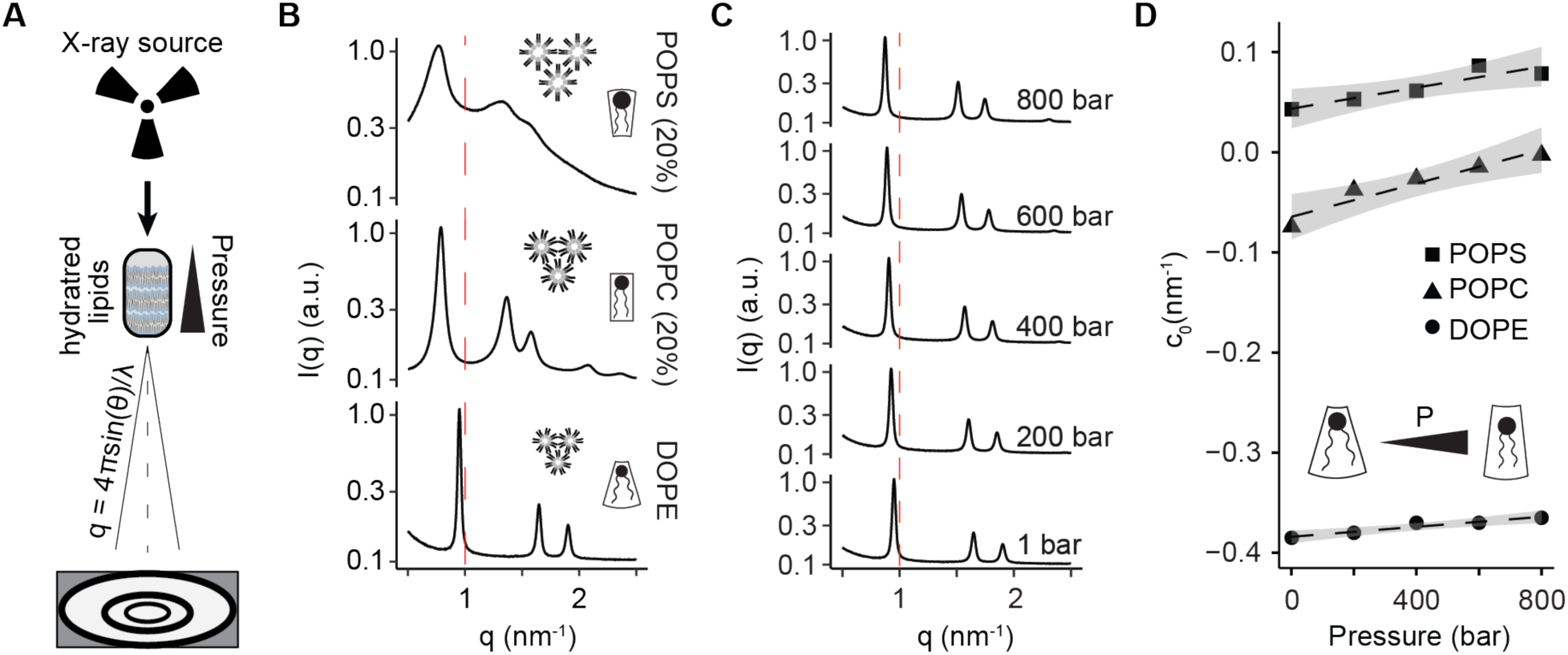
Hydrostatic pressure increases phospholipid intrinsic curvature. **A**) A schematic of an X-ray scattering setup where the scattering pattern on the detector can be used to determine the structural properties of lipid phases, including spontaneous curvature (c0), from their corresponding scattering intensity profiles. Hydrostatic pressure can be applied to hydrated lipid films in a specialized sample cell to study the effects of pressure on lipids. **B**) Bragg peak profiles of 100% DOPE or 20 mol% POPS or POPC hosted in 80 mol% DOPE, each with 12% w/w tricosene. The q-spacing of the peaks reflect the inverted hexagonal phase (HII) and the position of the first order peak is related to the lattice d-spacing and total spontaneous curvature. Addition of POPS and POPS results in leftward shifting of the first order peak in q-space. **C**) Bragg peak profiles of DOPE under 1-800 bar hydrostatic pressure. As pressure increases, the first order peak of the HII phase shifts leftward in q-space, representing a reduction in negative curvature. **D**) Spontaneous curvature values of c0 for DOPE, POPC, and POPS measured by HPSAXS across 1-800 bar hydrostatic pressure at 35 °C. As pressure increases, lipid negative curvature decreases by a similar slope in all 3 lipid classes.

### Lipid intrinsic curvature predicts hemifusion rates across variations in PE/PC and pressure

The above results suggest that both pressure and PE content affect fusion through their common influence on lipid intrinsic curvature. To test this directly, we calculated the corrected mean spontaneous curvature c̄_0_ for each of the 12 experimental conditions (3 compositions x 4 pressures) using the independently measured phospholipid curvatures at different pressures with HPSAXS (Fig. 1D). The c̄_0_ was computed as a weighted sum of the mole fraction and pressure-corrected intrinsic curvature of each lipid species. The resulting c̄_0_ ranged from −0.175 nm^−1^ (high PE at 1 bar) to −0.060 nm^−1^ (low PE at 750 bar) (Fig. 4A).

When rate constants from all 12 conditions were plotted against the corrected mean spontaneous curvature, the data collapsed in a semilogarithmic plot of ln(k) versus c̄_0_ spanning nearly two orders of magnitude in rate with a linear relationship (Fig. 4B). Data points from different compositions and different pressures fell along the same relationship, suggesting that mean intrinsic curvature governs the hemifusion rate regardless of whether it was varied by changing the PE:PC ratio or by applying hydrostatic pressure.

**Figure 4:**
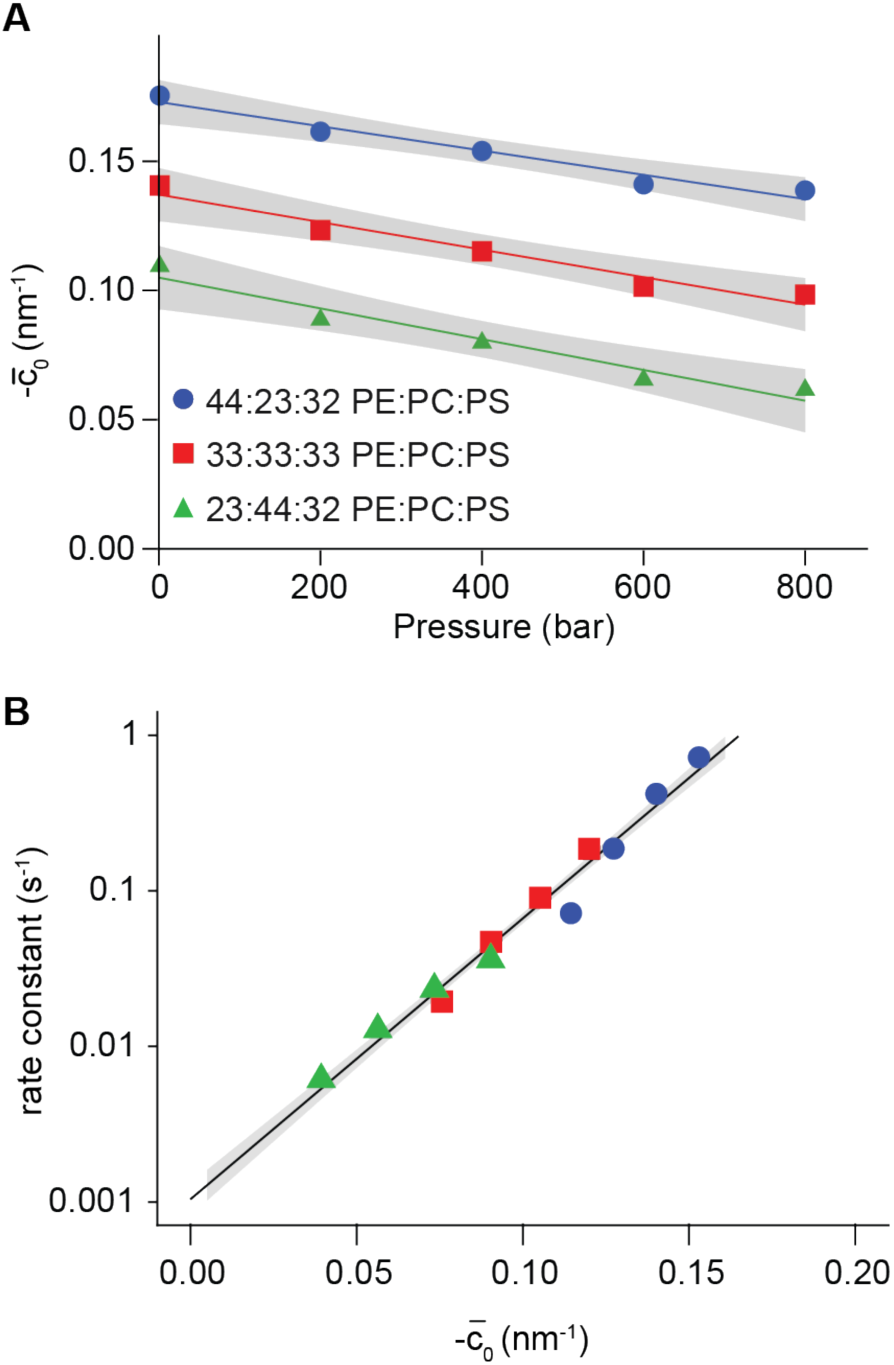
Hemifusion rate has an exponential dependence on mean lipid intrinsic curvature of liposome mixtures. **A**) Estimations of mean lipid intrinsic curvatures for 44:23:32 mol%, 33:33:33 mol%, or 23:44:32 mol% DOPE:POPCPOPS liposome compositions across 1-800 bar pressure. Mean curvature is calculated as a weighted sum of products of the mol% lipid and its c0 measured over a range of pressures by HPSAXS. SEM for the fit is shown in grey. **B**) The effect of corrected mean curvature on lipid mixing rate from 44:23:32 mol%, 33:33:33 mol%, or 23:44:32 mol% DOPE:POPC:POPS liposome compositions under 1-750 bar pressure shown on semilog axes. Error bars indicate SEM from n=3 replicate runs.

### Quantifying the relationship between c̄_0_ and the hemifusion energy barrier

To understand the relationship between curvature and hemifusion rate in energetic terms, we applied Kramers’ rate principle, which states that the rate of crossing an energy barrier increases exponentially with the height of that barrier. By this principle, the ratio of hemifusion rates, τ_1_ and τ_0_, at two different conditions reports the change in barrier energy between them, without requiring knowledge of the barrier height or the molecular attempt frequency. Previous studies have identified the hemifusion stalk as the key high-energy intermediate controlling the kinetics of lipid mixing (1). To calculate the change in this energy as a function of c̄_0_ we used continuum tilt-splay elastic model (7, 33), which expresses the monolayer energy density as:

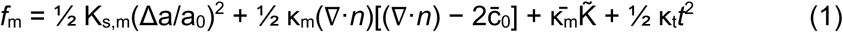

K_s,m,_ κ_m_, K^−^_m_ and κ_t_ are the monolayer stretching splay, saddle-splay, and tilt moduli, respectively. Δa/a_0_ denotes the local relative area change, ∇·*n* is the lipid splay, K̃ is the saddle-splay and *t* tilt. The overall stalk energy is obtained by integrating this energy density over both monolayers of the stalk and subtracting the energy of the pre-fusion planar bilayers:

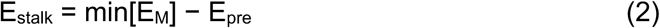

The hemifusion time rate ratio, ln(τ_1_/τ_0_), is proportional to the change in stalk energy as a function of pressure. We postulate that the primary effect of pressure is to alter c̄_0_, while the elastic moduli have negligible pressure dependence over the range considered, as supported by the modest change in lamellar spacing under pressure (Supplementary Fig. 1A-B). Under this assumption, all terms except the splay cross-term **∇**·***n*** cancel, and the change in barrier energy simplifies to:

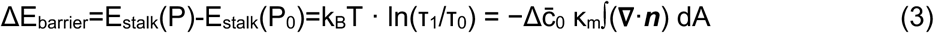

The slope of ΔE_barrier_ versus Δc̄_0_, κ_m_∫(**∇**·***n***)dA, quantifies the sensitivity of the fusion barrier to lipid curvature and is a property of the transition-state geometry, independent of c̄_0_.

Converting our measured rate constant ratios into barrier energy changes ΔE and plotting against Δc̄_0_, the energy barrier for hemifusion increased across the full range of curvatures sampled (colored lines, Fig. 5A). Between compositions, which separate pressure induced changes in c_0_ from confounding variations in elastic moduli, slopes ranged from 20.8 kT·nm to 31.5 kT·nm. The smaller values corresponded to the low-PE mixtures, suggesting a modest effect of lipid composition on stalk energy independent of c_0_. The pooled slope corresponds to a relationship between curvature and hemifusion energy of 32.6 ± 2.4 kT·nm.

To compare our results with the theoretical stalk model, we simulated stalk structures and calculated the splay integral over the monolayer surface, ∫(**∇**·***n***)dA, as a function of Δc̄_0_. The splay distribution depends on the monolayer thickness (Supplementary Fig. 2A), the ratio of the saddle-splay modulus to the splay modulus, *χ* (Supplementary Fig. 2B), and the tilt decay length, *l*, which is determined by the ratio of the tilt modulus to the splay modulus (Supplementary Fig. 2C). We evaluated the stalk structure over the relevant parameter range, considering monolayer thicknesses of 1–2 nm, tilt decay lengths of 1–2 nm, and values of *χ* between -1 and 0. Across this range, the splay integral varied between 12 and 22 nm. For representative parameter values *χ*=-1, l=1.0, and a monolayer thickness of 1.5 nm, the splay integral is approximately 16 nm. Fig. 5B shows a representative example of the stalk energy as a function of Δc̄_0_ for different values of the tilt decay length, *l*. The inset depicts the corresponding simulated stalk structure.

Using this range of splay integrals, we estimate that κ_m_ required to reproduce the experimentally inferred slope is 3-5 k_B_T (full black lines, Fig. 5A) This range is consistent with the membrane bending rigidities reported in the literature for DOPE:POPC:POPS membranes (19), which are twofold larger than the corresponding monolayer bending rigidities (34). Taken together, the estimated splay integral for the barrier configuration is in quantitative agreement with that predicted by the stalk model, supporting the interpretation of the barrier state as a stalk-like intermediate.

**Figure 5:**
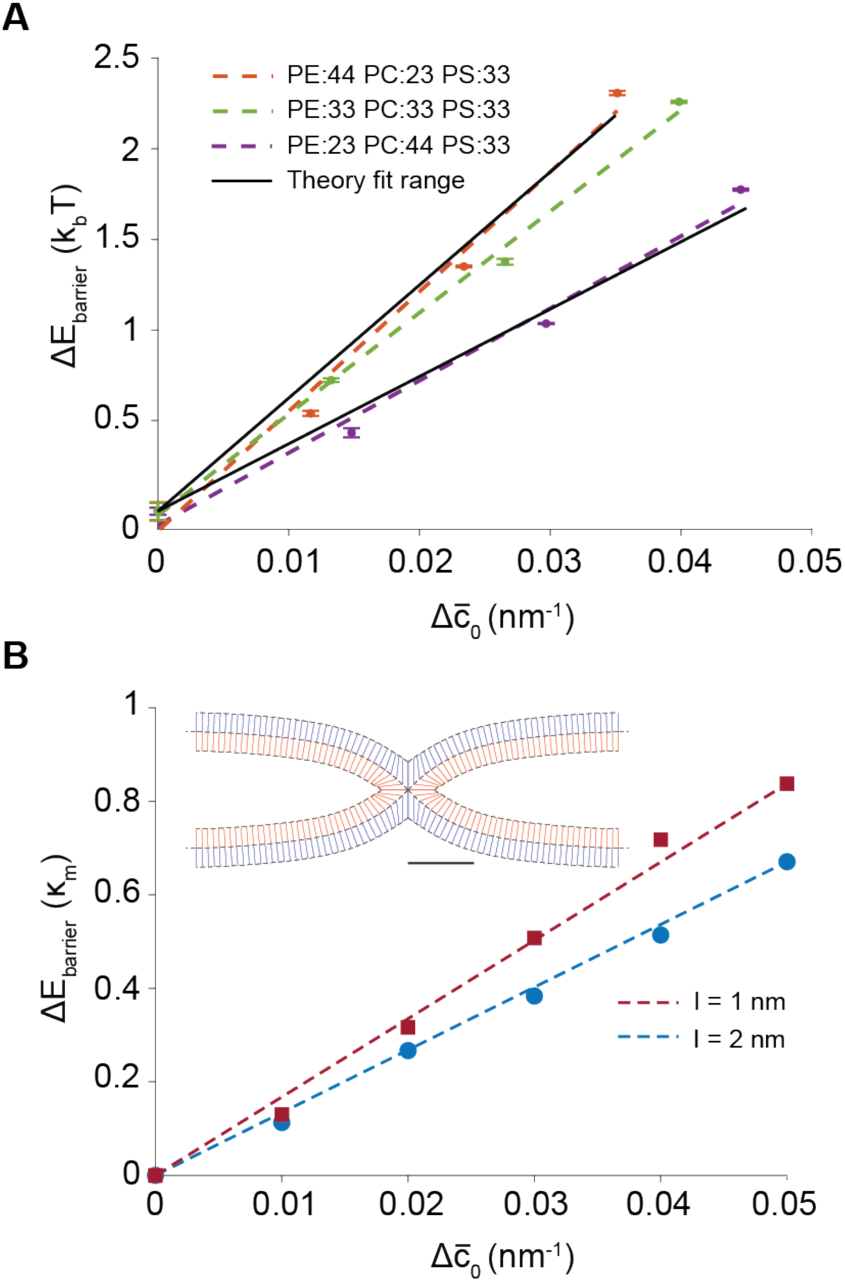
Stalk formation energies as function of. Δc̄0**. A**) Change in the stalk energy barrier with change in Δc̄0 for 44:23:32 mol%, 33:33:33 mol%, or 23:44:32 mol% DOPE:POPC:POPS liposome compositions as derived from Eq. 3. Error bars indicate SEM from n=3 replicates. The black lines represent fitting to the stalk model with *χ*=-1, l=1.0, and a monolayer thickness of 1.5 nm. The upper line corresponds to κm 5 kBT and the lower to 3 kBT. **B)** Representative example of the change in stalk energy as a function of Δc̄0, normalized by the monolayer bending rigidity κm. Simulations were performed for a monolayer thickness of 1.5 nm and χ=-1. The two curves correspond to the lower and upper bounds of the tilt decay length, *l*, reported in the literature. Inset: Simulation snapshot for a tilt decay length of 1.5 nm and c̄0=0. Scale bar, 5 nm

## Discussion

The linear relationship between lipid intrinsic curvature and hemifusion stalk formation energy has been a central prediction of the stalk hypothesis, yet it has not previously been tested in a system where c_0_ is varied independently of lipid chemistry. By combining hydrostatic pressure with independent HPSAXS calibration of each lipid species, we isolated c_0_ as a single variable and found that all 12 experimental conditions, spanning three lipid compositions and four pressures, collapse onto a single linear ln(k) versus c̄_0_ relationship. The corrected c̄_0_ appears to be the primary variable governing the hemifusion rate regardless of whether it was varied by changing the PE:PC ratio or by applying pressure.

The curvature–energy slope from our pressure measurements (20–30 k_B_T·nm) is consistent with values obtained from a wide range of experimental fusion systems. Based on the reported effects of lipid composition on hemifusion rates, and using published estimates of the c_0_ of the perturbed lipids to convert the compositional changes into changes in monolayer spontaneous curvature (35), we estimate slopes of ∼25 k_B_T·nm and ∼10 k_B_T·nm for hemagglutinin and SNARE-mediated fusion, respectively, using cholesterol perturbations (12, 13) and ∼6–20 k_B_T·nm for protein-free Ca^2+^-mediated measurements using cholesterol and LPC perturbations (14). These values are not reported directly in the cited works but were derived under the assumptions that compositional perturbations primarily affect intrinsic curvature rather than elastic moduli, that the intrinsic curvatures of the individual lipids are known, and that monolayer spontaneous curvature can be approximated as a composition-weighted average. Despite the crude nature of these estimations, the agreement among measurements obtained from protein-free, SNARE-mediated, and viral-fusion systems, as well as from distinct experimental approaches, suggests that the effect of spontaneous curvature on the hemifusion energy barrier is constrained to the range of approximately 10–30 k_B_T·nm.

The experimentally inferred range of the hemifusion energy barrier is substantially lower than predictions from earlier continuum elastic models and molecular dynamics simulations (∼200-300 k_B_T·nm) (7, 8, 10, 11). One possible source of this discrepancy is that earlier continuum models imposed restrictive geometric assumptions on the stalk structure. In the present work, we refined the elastic model by allowing the junction angle to vary freely and adopt the value that minimizes the total elastic energy. This additional degree of freedom relaxes the local splay stress and substantially reduces the sensitivity of the stalk energy to spontaneous curvature. Furthermore, the splay distribution within the stalk depends strongly on other elastic parameters, particularly the tilt modulus through the tilt decay length, *l*, whose value remains poorly constrained experimentally. Consequently, estimates of the dependence of stalk energy on curvature are sensitive to assumptions regarding other elastic moduli.

Within this refined framework, the stalk model can be reconciled with experimental observations if the monolayer splay modulus is relatively soft, approximately 3–5 k_B_T, and the tilt decay length is short, on the order of 1 nm or less. Importantly, such values do not necessarily imply unusually soft bilayers. The monolayer bending modulus used in many previous models was inferred from the common assumption that the bilayer bending modulus is simply twice the monolayer value. However, this relation is valid only when lipids can freely exchange between the two leaflets during membrane deformation. If interleaflet lipid exchange is slower than the characteristic deformation timescale, additional area-compression stresses develop and contribute to the measured bilayer bending modulus (36). Under these conditions, the true monolayer bending modulus can be substantially lower than half the bilayer value, yielding a softer monolayer that is nevertheless fully consistent with experimentally measured bilayer elastic properties and with the curvature sensitivity of stalk energy measured here. Resolving this discrepancy further will require independent experimental constraints on the monolayer bending modulus and tilt decay length, potentially through neutron spin echo measurements of monolayer fluctuation dynamics or through single-leaflet mechanical probes that bypass the ambiguity of the bilayer-to-monolayer conversion.

Beyond testing the stalk hypothesis, the effect of c̄_0_ of fusion provides quantitative insight for understanding how lipid composition tunes fusion kinetics in living cells. Eukaryotic membranes maintain approximately 30–40% negatively curved lipids including PE, cardiolipin, and DAG, positioning the average membrane curvature in a narrow window of c̄_0_ ≈ −0.05 to −0.15 nm^−1^ (6, 37). At a slope of ∼32 kT·nm, shifting c̄_0_ by 0.01 nm^−1^, equivalent to replacing roughly 5–10 mol% PC with PE, would alter the fusion rate by approximately 1.5-fold. These effects are biologically meaningful, particularly in contexts where fusion is precisely timed such as synaptic exocytosis, yet moderate enough to buffer against stochastic lipid composition fluctuations. Cells appear to exploit this sensitivity through enzymatic pathways that locally remodel lipid curvature at sites of membrane trafficking, including phospholipase D generating phosphatidic acid (PA) at exocytic sites (38), DAG kinases interconverting DAG and PA (39), and PE methyltransferases converting PE to PC (40). The localization of these enzymes to membranes where fusion or fission occurs suggests that local lipid curvature remodeling serves to tune the kinetic competence of membranes for topological rearrangement. Studies have suggested that mammalian cells actively maintain intrinsic curvature stress on the lipidome level (41, 42), analogous to homeoviscous adaptation for membrane fluidity (43).

Our results also provide evidence that hydrostatic pressure inhibits membrane hemifusion through its effect on lipid intrinsic curvature, connecting *in vitro* biophysics to emerging observations of pressure adaptation in living organisms. Deep-sea ctenophores increase their abundance of plasmalogen PE, a lipid with strongly negative c_0_, in proportion to their habitat depth (44), and yeast with reduced PE/PC ratios show impaired growth and viability at elevated pressure independently of membrane fluidity (42). In both cases, high pressure diminishes the accessibility of nonlamellar lipid phases, which has been reasoned to compromise membrane fusion and cellular fitness (45). Our data provide the mechanistic basis for these observations by showing that the pressure-induced shift in c_0_ slows hemifusion stalk formation. The approximately 10-fold reduction in rate constant across 750 bar at equimolar PE:PC:PS composition, and the ability of elevated PE content to buffer against this inhibition, are consistent with the organismal strategies observed in pressure-adapted species. These findings suggest that the evolutionary pressure to maintain negatively curved lipids in biological membranes reflects a requirement to keep the hemifusion barrier within a range accessible to the cell’s fusion machinery.

Several limitations of this study should be considered. Our model system uses Ca²⁺/PS-mediated fusion of LUVs, which lacks the protein machinery that drives fusion *in vivo*. While this simplification allows us to isolate lipid contributions to hemifusion without confounding pressure effects on proteins, it remains to be determined whether the measured relationship between curvature and stalk energy holds in protein-mediated systems where fusogens impose additional geometric constraints on the stalk. The three-component lipid mixtures used here do not capture the complexity of biological membranes, which contain hundreds of species that contribute to curvature stress and elastic properties. Moreover, ternary mixtures near demixing boundaries can exhibit mechanical softening that dampens bending and other elastic moduli, potentially influencing the measured sensitivity of stalk energy to curvature independently of mean spontaneous curvature (46). Although our compositions are not expected to lie near a demixing transition at 37 °C, proximity to such a boundary cannot be ruled out in the presence of Ca²⁺, which can induce PS clustering. Our calculation of mean spontaneous curvature also assumes ideal mixing and a weighted average, neglecting potential demixing and compositional heterogeneity at the fusion site. Local regions of high curvature may promote selective sorting of lipids like PE. Additionally, our FRET-based assay reports lipid mixing as a proxy for hemifusion but does not report on fusion pore formation, which is governed by distinct energetic considerations. Finally, our analysis assumes that pressure-dependent changes in elastic moduli are negligible over the range studied. While the small pressure sensitivity of bilayer thickness (Supplementary Fig. 2) supports this assumption, the saddle-splay (Gaussian) modulus, which contributes to stalk energy through the topological change inherent in hemifusion, remains poorly constrained for most lipid compositions and has been challenging to measure experimentally (47).

## Conclusions

By exploiting the unique ability of hydrostatic pressure to tune lipid intrinsic curvature without altering lipid chemistry, we have provided experimental support for the predicted linear relationship between c_0_ and the hemifusion stalk formation energy. The collapse of data from multiple compositions and pressures onto a single master curve suggests that the corrected mean spontaneous curvature is the dominant variable controlling hemifusion rate in this system. The measured slope of 20–30 kBT·nm supports predictions of the stalk hypothesis when considered alongside refined continuum elastic models with realistic monolayer bending moduli. More broadly, the approach demonstrated here, utilizing hydrostatic pressure to manipulate lipid geometry, provides a general strategy for dissecting the individual contributions of biophysical properties to membrane remodeling processes.

## Acknowledgements

Raya Sorkin and Michael Kozlov provided helpful discussions and feedback. Cathy Royer and the Royer lab provided technical support for high-pressure stopped flow fluorometry. Research was supported by the National Science Foundation (NSF) (MCB-2046303 and MCB-2316458 to I.B.), the National Institutes of Health (5T32EB009380-14 to D.M.), NASA (Postdoctoral fellowship 0017-NPP-MAR22-A-Astrobio to J.R.W.) and Israel Science Foundation (1367/25 to G.G). The High-Pressure Stopped Flow System was supported by the NSF-MRI Award Number (FAIN): 2213116.

## Author contributions

D.M., J.R.W, G.G, and I.B conceived the project and designed the analyses. D.M, J.R.W, and J.E.M. carried out experiments. G.G. carried out modeling. D.M. and G.G. wrote the manuscript. D.M., J.R.W, G.G., and I.B. revised the manuscript. G.G. and I.B. acquired funding.

## Supplementary Information

**Supplementary Figure 1:**
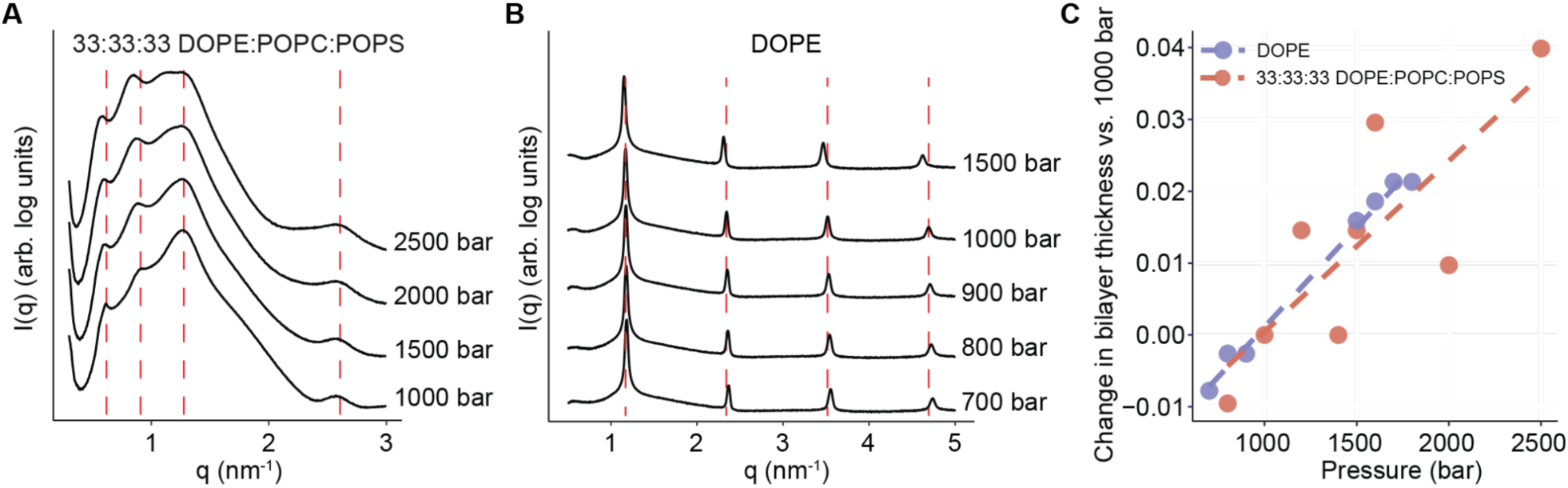
Change in lamellar spacing of model membranes under pressure. **A**) SAXS scattering profiles of a 33:33:33 DOPE:POPC:POPS lipid mixture in 10 mM CaCl_2_ buffer at 35 °C and 1000-2500 bar pressure. Vertical lines indicate peak positions at 1000 bar. **B**) SAXS scattering profiles of pure DOPE in water at 35 °C and 700-1500 bar pressure. Vertical lines indicate peak positions at 1000 bar. **C**) Change in bilayer thickness from a 1000 bar reference pressure for 33:33:33 DOPE:POPC:POPS lipid mixture in 10mM CaCl_2_ buffer and DOPE in water at 35 °C.

**Supplementary Figure 2:**
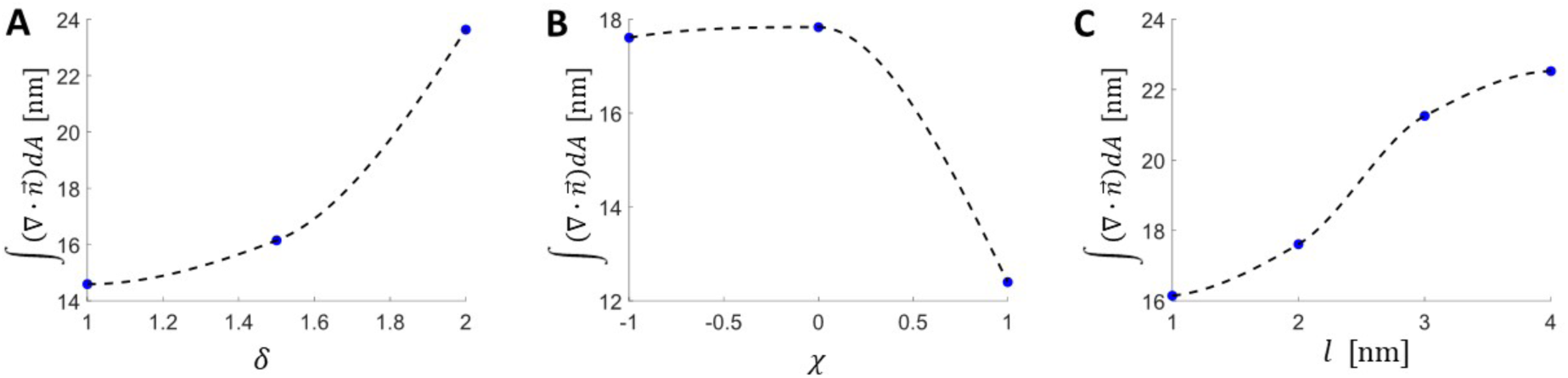
Splay integral, ∫(**∇**·***n***)dA, as a function of different parameters obtained from numerical calculations of stalk structure. **A**) as a function of the distance between the membrane mid-plane to the monolayer neutral plane δ. χ=-1 and *l*=1 nm. **B**) as a function of the ratio between monolayer saddle-splay to splay modulus (χ=K^−^_m_/κ_m_). δ=1.5 nm and *l*=2 nm. **C**) as a function of the tilt decay length, 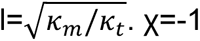 and δ=1.5 nm.

